# Pharmacological AMP-activated protein kinase activation suppresses low glucose-dependent macrophage migration inhibitory factor release from macrophages

**DOI:** 10.1101/2023.10.16.562445

**Authors:** Jiping Zhang, Alice E. Pollard, David Carling, Benoit Viollet, Kate L.J. Ellacott, Craig Beall

**Author notes:** Corresponding Author: Dr Craig Beall; Postal address: RILD Building, Level 4, Room 4.06, Barrack Road, Exeter, EX2 5DW Email address.

## Abstract

**Aims/hypothesis:** Acute hypoglycemia promotes pro-inflammatory cytokine production, increasing risk for cardiovascular events in diabetes. AMP-activated protein kinase (AMPK) is regulated by and influences production of pro-inflammatory cytokines. We tested the mechanistic role of AMPK in low glucose induced changes in the pro-inflammatory cytokine macrophage migration inhibitory factor (MIF), which is elevated in patients with diabetes.

**Methods:** Macrophage cell line Raw264.7 cells, primary macrophage bone marrow derived macrophages obtained from wild type mice or AMPK γ1 gain-of-function mice were utilized, as were AMPKα1/α2 knockout mouse embryonic fibroblasts (MEF). Allosteric AMPK activators PF-06409577 and BI-9774 were used, in conjunction with inhibitor SBI-0206965 were also used. We examined changes in protein phosphorylation/expression using western blotting, and protein localisation using immunofluorescence. Metabolic function was assessed using extracellular flux analyses and luciferase-based ATP assay. Cytokine release was quantified by ELISA. Oxidative stress was detected using a fluorescence-based ROS assay, and cell viability was examined using flow cytometry.

**Results:** Macrophages exposed to low glucose showed a transient and modest activation of AMPK and a metabolic shift towards increased oxidative phosphorylation. Low glucose induced oxidative stress and increased release of macrophage migration inhibitory factor (MIF). Pharmacological activation of AMPK by PF-06409577 and BI-9774 attenuated low glucose-induced MIF release, with a similar trend noted with genetic activation using AMPKγ1 gain-of-function (D316A) mice, which produced a mild effect on low glucose-induced MIF release. Inhibition of NFĸB signalling diminished MIF release and AMPK activation modestly but significantly reduced low glucose-induced nuclear translocation of NFĸB. AMPK activation did not alter low glucose-induced oxidative stress in macrophages but application of AMPK inhibitor SBI-0206965 enhanced oxidative stress in macrophages and in AMPK knockout MEFs, suggesting an AMPK-independent mechanism

**Conclusions/interpretation:** Taken together, these data indicate that pharmacological AMPK activation suppresses release of MIF from macrophages. This is mediated by reduced activation of NFĸB signalling in response to low glucose-induced oxidative stress and suggests that pharmacological AMPK activation could be a useful strategy for mitigating hypoglycemia-induced inflammation.

**Graphical abstract:** Tweet
Low glucose induces pro-inflammatory MIF release from macrophages, and pharmacological AMPK activation suppresses the release of MIF. AMPK/ NFĸB signalling pathway is involved, which may be a new strategy to attenuate the pro-inflammatory response in hypoglycemia.

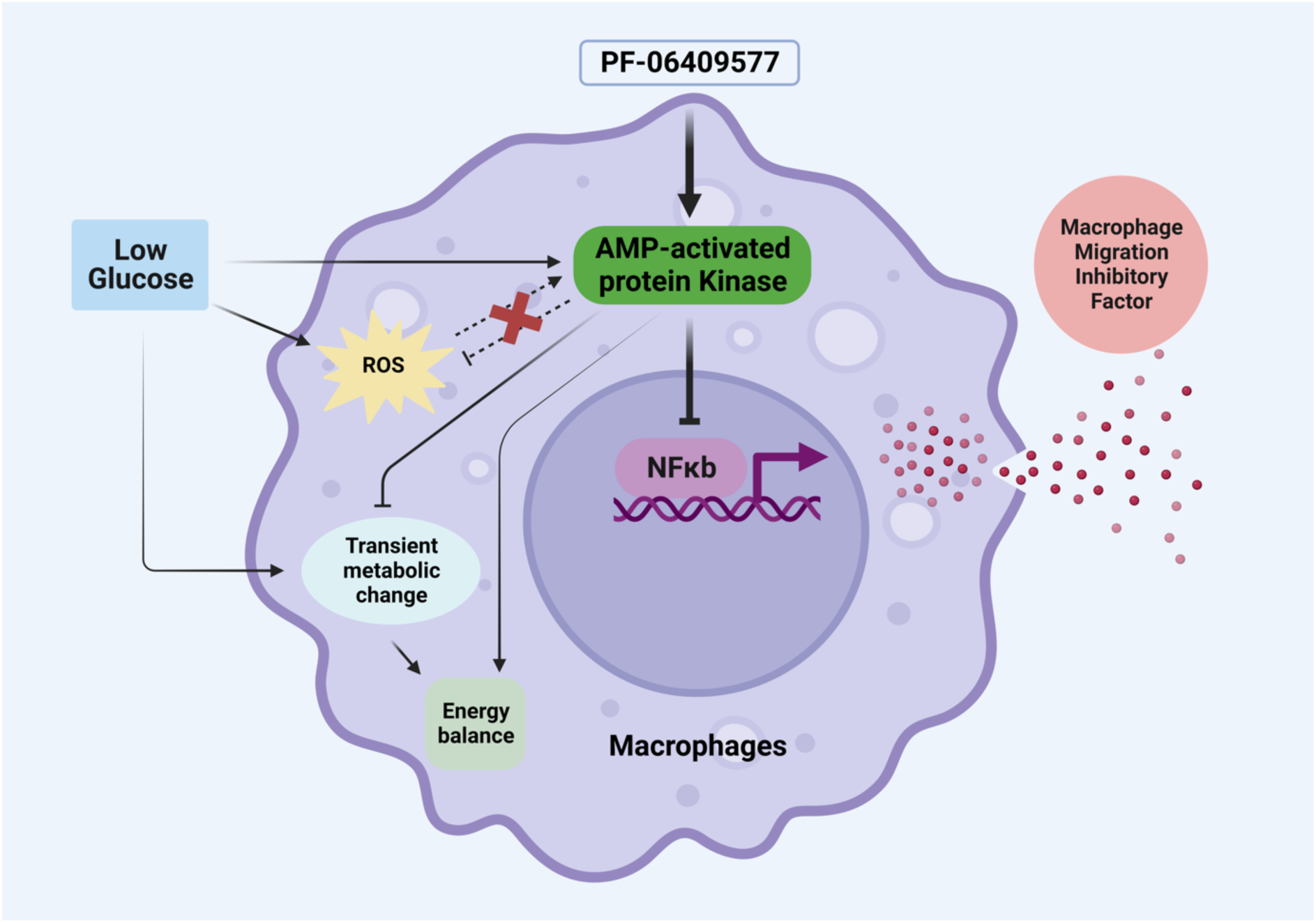

## 1. Introduction

Chronic low-grade inflammation contributes to the pathogenesis of diabetes [3]]. People with diabetes have higher circulating levels of inflammatory markers including macrophage migration inhibitor (MIF) [4], C-reactive protein (CRP), interleukin-6 (IL-6) and tumour necrosis factor-α (TNF-α) [5]. MIF is a cytokine expressed broadly in the outer lining of tissues such as endothelial, epithelial, endocrine and immune cells including macrophages and can be released in response to cellular stress from readily releasable pools [6].Plasma MIF levels are elevated in patients with impaired glucose tolerance and further increased in Type 2 diabetes [4, 7, 8]. MIF is also highly expressed in diabetes complications such as myocardial damage [9], coronary artery disease and diabetic retinopathy [8], suggesting a graded increase in circulating MIF with disease severity. The plasma levels of MIF also correlate with the severity of illness in COVID-19 patients [10]. The cytokine increases production of IL-1β, IL-6 and TNFα [6] and may also play a role in thrombosis [11]. In contrast, MIF deficiency [12] or antagonism [13] has been shown to prevent chronic inflammation, glucose intolerance and insulin resistance in mice, suggesting MIF likely plays a detrimental role in diabetes.

Recent studies have demonstrated that hypoglycemia in addition to hyperglycemia contributes to inflammation via increasing circulating markers of inflammation such as C-reactive protein (CRP) and oxidative stress [14]. Correlating to MIF-induced changes in cytokines, hypoglycemia also increases the levels of IL-1β, IL-6 and TNFα [15]. Importantly, moderate and severe hypoglycemia increase risk for a cardiovascular event [16] and is also associated with neurological damage and increased risk for mortality [17]. As a major source of cytokines including MIF, macrophages play an important role in chronic inflammation in diabetes. This response is an energy-intense process, with macrophage metabolism shifting from mainly oxidative metabolism to aerobic glycolysis during the pro-inflammatory phase [18]. This switch is associated with reduced activation of the key energy sensor 5ʹ adenosine monophosphate-activated protein kinase (AMPK) activation. This metabolic switch in macrophages can be inhibited by pharmacological activation of AMPK [19]. The kinase senses ratios of ATP to AMP and ADP and restores energy balance by inhibiting anabolic processes that consume ATP, while promoting catabolic processes that generate ATP [20]. AMPK also plays an anti-inflammation role. The first-line Type 2 diabetes drug metformin, which also activates AMPK [21], reduces systemic inflammation by decreasing the level of CRP and IL-6 [22] and lowers plasma macrophage migration in obese patients [19]. Together these data suggest that AMPK is an attractive drug target to improve metabolic homeostasis and reduce inflammation. Here we used pharmacological AMPK activators, PF-06409577 [23] given its use in a recent phase I clinical study (NCT02286882) and BI-9774, in conjunction with AMPK gain-of-function mice, to examine whether AMPK activation could prevent or reduce pro-inflammatory MIF production from macrophages following energy stress.

## 2. Materials and Methods

### 2.1 Cell isolation and culture

Raw264.7 cells and primary BMDMs were cultured as previously described [24]. Bone marrow was collected from C57BL/6J wild-type healthy mice. Immediately before treatments, cells were cultured in serum-free DMEM or RPMI medium with 5.5 mmol/l glucose then transferred to experimental conditions. AMPKγ1 gain-of-function mice (*prkag1*, aspartic acid to alanine at residue 316; [D316]) were generated as described previously [25] and BMDMs isolated from male wild type (WT) or AMPKγ1 D316A mutant mice. Mouse embryonic fibroblasts (MEF) WT and AMPK α1/2 (-/-) double knock-out cells were generated as previously described [26] and were cultured as described before [27].

### 2.2 Animals

Mice (male) were housed in pathogen-free individually ventilated cages on a 12hr light/dark cycles and maintained at a constant temperature of 22±1°C. Mice were group housed (up to 4 per cage) and *ab libitum* fed a standard diet (Exeter: 2018, Teklad Global; Imperial: RM3 diet, Special Diet Services;) with free access to water. Animals were provided woodchip bedding, tissue nesting paper with environmental enrichment (tunnels and chew sticks). Animals were culled by Schedule 1 method and all studies were performed in accordance with the United Kingdom Animals (Specific Procedures) Act (1986) and approved by the Animal Welfare and Ethical Review Boards at the University of Exeter and Imperial College London.

### 2.3 Western blotting

Cellular protein was isolated as previously described [28]. Briefly, 2.5×10^6^ Raw264.7 cells were seeded onto 60 mm petri dishes one day before treatment and harvested lysis buffer. BMDMs were defrosted from liquid nitrogen (4-7 × 10^6^ BMDMs/dish) were seeded onto 60 mm petri dishes one day before treatment. Protein concentrations were assessed by the method of Bradford. 10 μg protein of each sample was loaded on either 7%, 10% or 15% (wt/vol.) polyacrylamide gels. Proteins were transferred to nitrocellulose membranes, blocked with powdered milk (5% wt/vol.) in TBS/T and probed with antibodies against target proteins followed by appropriated tagged secondary antibodies. Images were taken using infrared imaging and changes in expression quantified by densitometry. Changes in phosphorylation were normalized to total expression of the target protein.

### 2.4 Immunocytochemistry

Immunocytochemical staining were performed as previously described [29]. Briefly, cells were plated on 13 mm diameter glass coverslips. After treatment, cells were fixed, permeabilized, non-specific binding blocked using donkey serum before incubating with mouse anti-NFĸBp65 antibody (1:1000 in TBST; overnight at 4 °C) and secondary donkey anti-mouse Alexa fluor488 (1:500 in PBS-T) for 1 hour at room temperature. Coverslips were mounted in DAPI-containing mounting media (ab104139, Abcam, Cambridge, UK) before imaging using a Leica SP8 confocal microscope. FIJI software was used to manually analyse fluorescent staining.

### 2.5 Enzyme-linked immunosorbent assay (ELISA)

Cytokines secreted into culture supernatants were measured by ELISA according to the manufacturers’ protocols (DY1978, Biotechne, Abingdon, UK). Briefly, media supernatants were collected and diluted, as required. Extracellular cytokine levels were normalized to total protein, unless otherwise stated.

### 2.6 Measurement of extracellular and intracellular ATP levels

ATP levels were measured using ATPlite assay kit (6016941, Perkin Elmer, Seer Green, UK) as previously described [27]. For extracellular ATP, 100 µl of media supernatant was used per sample, in black-walled 96 well plates. The luminescence was read using the PheraStar microplate reader (BMG LabTech). ATP concentrations were calculated and normalized to total protein. Raw264.7 cells were plated in black-walled 96 well plates 24 hours before treatments then cells were lysed and luminescence was read and intracellular ATP levels was presented as pmol/l.

### 2.7 Measurement of cellular metabolism and viability

Raw 264.7 cells and BMDMs were seeded in Seahorse XF2 96 culture plates (102416-100, Agilent, Craven Arms, UK) 24 hours prior to treatment. Metabolic measurements were performed following the manufacturer’s instructions. Briefly medium was exchanged with DMEM or RPMI Seahorse XF medium pH7.4 and cells degassed for 1hr. Extracellular acidification rate (ECAR) as a measure of glycolysis, and oxygen consumption rate (OCR) a measure of mitochondrial function normalized to the total protein content. Cell viability assay was performed by propidium iodide (PI) (P4864, Merck, Gillingham, UK) staining followed by flowcytometry analysis [29]. See ESM for further details.

### 2.8 Reactive oxygen species (ROS) assay

Raw 264.7 cells were cultured in 48-well plates. Following treatments, medium was replaced with 10 µmol/l of DCFDA (D399; Invitrogen, Loughborough, UK) in DMEM and incubated at 37°C for 30 minutes. The levels of ROS were measured using PheraStar microplate reader. Relative fluorescence units were normalized to the total protein content.

### 2.9 Statistical analysis

Two-group comparisons were determined by t-test using GraphPad Prism software (Prism 9; GraphPad Software, La Jolla, CA, USA). For multiple group comparisons a one-way ANOVA with Bonferroni Statistical tests was performed. To compare the mean differences between groups split by two or three independent variables, a two-way ANOVA or a three-way ANOVA with Bonferroni multiple comparisons test was used respectively. Results are expressed as mean ± standard error. Values of p<0.05 were considered statistically significant.

## 3. Results

### 3.1 Physiological and pharmacological activation of AMPK in macrophages

AMPK is activated by reduced glucose availability in several cell types [28, 30]. To examine whether this also occurs in macrophages, we exposed Raw264.7 cells to normal (5.5 mmol/l) and low (1.0 mmol/l) glucose levels. Exposure to low glucose for 30 minutes increased phosphorylation of AMPK at threonine 172 (T172), a phosphorylation site required for full kinase activation (Fig. 1a). Phosphorylation of acetyl-CoA carboxylase (ACC, ser79), a downstream substrate of AMPK, was also increased (Fig. 1a), indicating increased AMPK activity. However, after 16 hours of low glucose, neither the level of AMPK or ACC phosphorylation was altered in either Raw264.7 cells or BMDMs (ESM Fig. 1).

**Figure 1.**
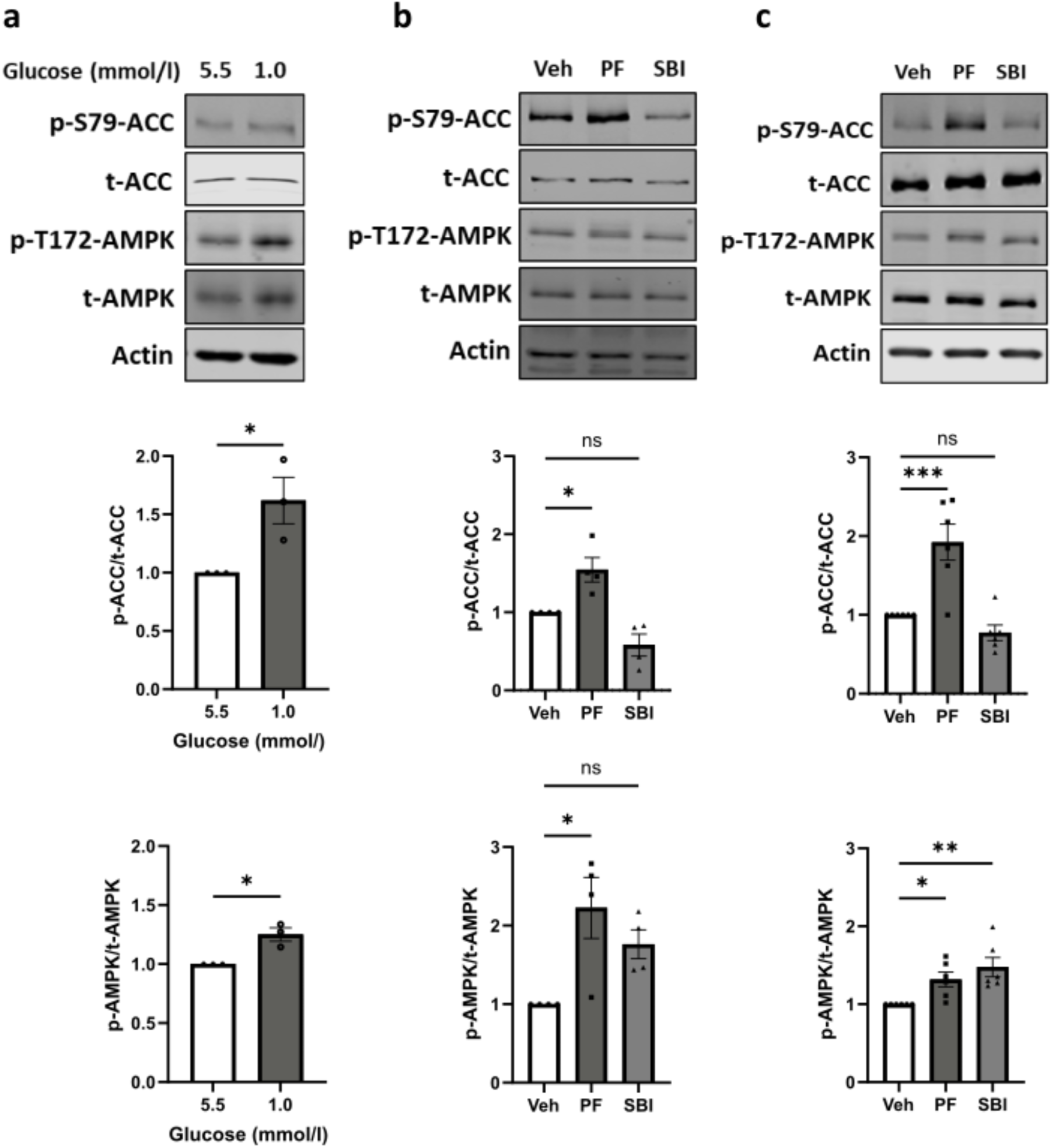
Low glucose and PF-06409577 increased phosphorylation of AMPK in macrophages. **a**) Representative immunoblots of Raw264.7 cells cultured with either 5.5 mmol/l or 1.0 mmol/l glucose for 30 minutes. **b**) Representative immunoblots of Raw264.7 exposed to vehicle (0.1% v/v DMSO) (Veh), 10 µmol/l of PF-06409577 (PF) or 30 µmol/l of SBI-0206965 (SBI) for 24 hours. **c**) Representative immunoblots of BMDMs exposed to vehicle (0.1% v/v DMSO) (Veh), 10 µmol/l of PF-06409577 (PF) or 30 µmol/l of SBI-0206965 (SBI) for 16 hours. Cells were then lysed and immunoblots were prepared. Densitometric analysis of immunostaining for phosphorylated protein was normalized to total protein level. p-ACC, phospho-acetyl-CoA carboxylase (S79) (p-S79-ACC); p-AMPK, phospho-AMP-activated protein kinase (T172) (p-T172-AMPK); t-ACC indicates total acetyl-CoA carboxylase; t-AMPK, total AMP-activated protein kinase. Data are expressed as mean±SEM (**a**, n=3; **b**, n=4; **c**, n=6). Comparisons between groups were made by unpaired t test (**a**) or one-way ANOVA with Bonferroni’s multiple comparisons test (**b**, **c**). *P<0.05; **P<0.01; ***P<0.001; ns, not significantly different.

We next examined the effect of pharmacological AMPK activation on macrophages by using PF-06409577. To confirm the effect of PF-06409577 and AMPK inhibitor SBI-0206965 on AMPK activation in macrophages, Raw264.7 and BMDMs were incubated in the presence of 10 µmol/l PF-06409577 or 30 µmol/l SBI-0206965 for 24 hours and 16 hours respectively. PF-06409577 increased phosphorylation of AMPK and ACC substantially in both macrophage cell types (Fig. 1b, c). Consistent with a previous report [2], SBI-0206965 increased AMPK phosphorylation at T172 yet showed a trend towards decreased pACC in both Raw264.7 cells and BMDMs (Fig. 1 b,c), suggesting AMPK inhibition, as expected.

### 3.2 Low glucose-induced MIF release from macrophages was attenuated by pharmacological AMPK activation

To examine the consequence of glucose variation on MIF release from macrophages, Raw264.7 cells were cultured in the presence of decreasing concentrations of glucose. Interestingly low glucose (1 mmol/l, 2.5 mmol/l, 3.5 mmol/l) induced MIF release in a concentration-responsive manner from Raw264.7 cells, compared with normal glucose (5.5 mmol/l; Fig. 2a,b). In contrast, high glucose (>10 mmol/l) did not alter MIF release from Raw264.7 cells (Fig. 2a). These differences could not be accounted for by a change in cell viability (ESM Fig. 3a)

**Figure 2.**
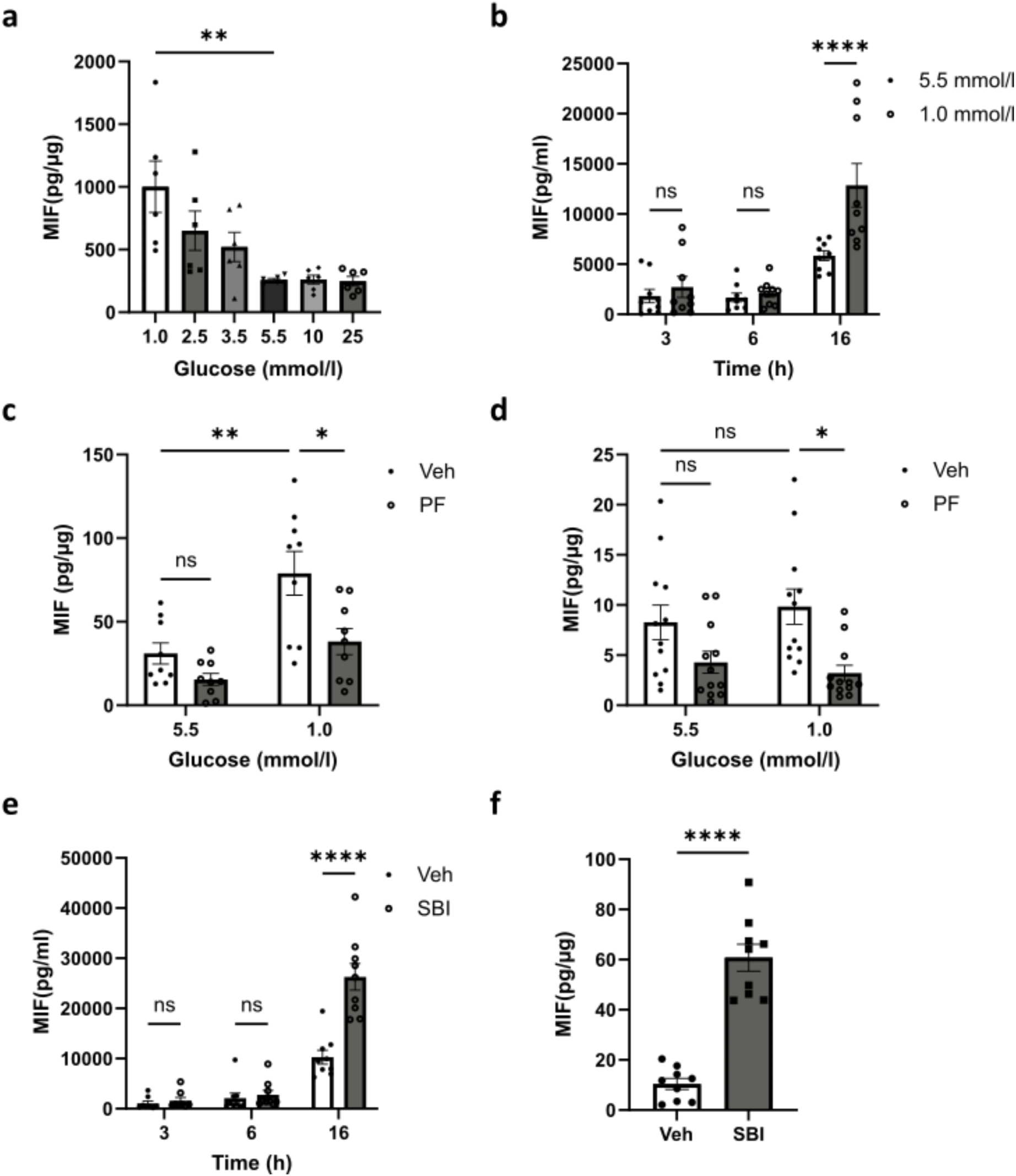
Low glucose (1.0 mmol/l) increased MIF release from Raw264.7 cells and MIF release was attenuated by PF-06409577 in Raw264.7 cells and BMDMs, but SBI-0206965 enhanced MIF release in both cell types. **a**) Raw264.7 cells were cultured with different concentrations of glucose (1.0, 2.5, 3.5, 5.5, 10 and 25 mmol/l) for 24 hours. **b**) Raw264.7 cells were cultured with normal glucose (5.5 mmol/l) or low glucose (1.0 mmol/l) for 3 hours, 6 hours and 16 hours. **c**) Raw264.7 cells and **d**) BMDMs were cultured with normal glucose (5.5 mmol/l) or low glucose (1.0 mmol/l) in the presence of vehicle (0.1% v/v DMSO) (Veh) or 10 µmol/l of PF-06409577 (PF) for 16 hours. **e**) Raw264.7 cells and **f**) BMDMs were cultured in the presence of vehicle (0.1% v/v DMSO) (Veh) or 30 µmol/l of SBI-0206965 (SBI) for 16 hours. The MIF level in medium was assessed by ELISA. The amount of released MIF was shown as concentrations (pmol/ml) (**b**, **e**), or normalized to the total protein amount in each sample (**a**, **c**, **d**, **f**). Data are expressed as mean±SEM (**a**, n=6; **b**, n=9; **c**, n=9; **d**, n=12; **e**, n=9; **f**, n=9). Comparisons between groups were made by unpaired t test (**f**), one-way ANOVA (**a**), or two-way ANOVA with Bonferroni’s multiple comparisons test (**b**, **c**, **d**, **e**). *P<0.05; **P<0.01; ****P<0.0001; ns, not significantly different.

**Figure 3.**
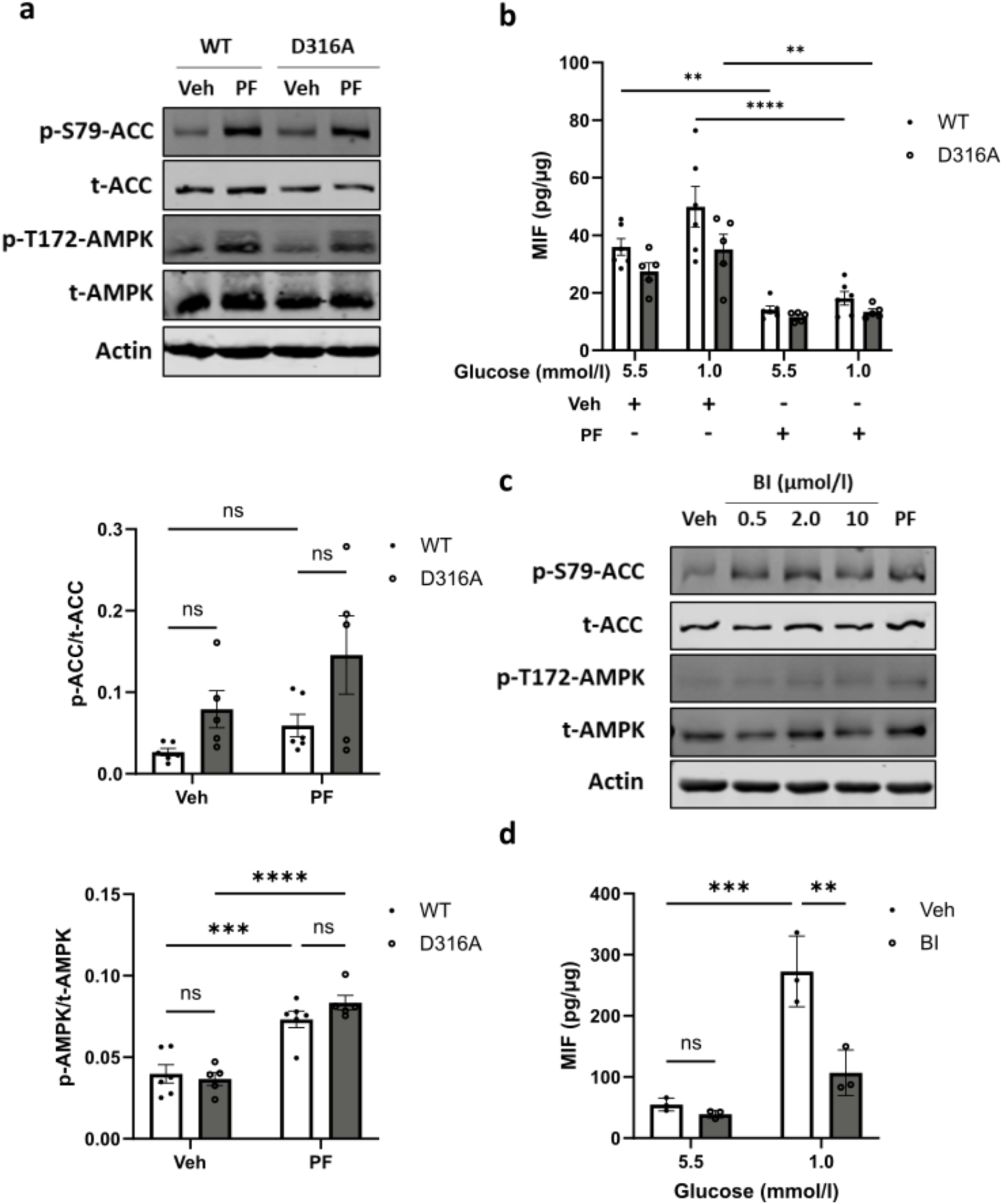
PF-06409577 activated AMPK and inhibited MIF release from both wild type and D316A BMDM and low glucose induced MIF release from Raw264.7 cells was attenuated by BI-9774. **a**) Representative immunoblots of BMDM exposed to vehicle (0.1% v/v DMSO) (Veh) or 10 µmol/l of PF-06409577 (PF) for 16 hours. BMDM cells were obtained from either wild type (WT) or AMPKγ1 D316A (D316A) mice. Densitometric analysis of immunostaining for phosphorylated protein was normalized to total protein level. P-ACC, phospho-acetyl-CoA carboxylase (S79) (p-S79-ACC); p-AMPK, phospho-AMP-activated protein kinase (T172) (p-T172-AMPK); t-ACC indicates total acetyl-CoA carboxylase; tAMPK, total AMP-activated protein kinase. **b**) BMDM cells obtained from either wild type (WT) or D316A mice (D316A) were cultured with normal level of glucose (5.5 mmol/l) or low level of glucose (1.0 mmol/l) in the presence of vehicle (0.1% v/v DMSO) (Veh) or PF-06409577 (PF) for 16 hours. The MIF release in medium was assessed by using ELISA. **c**) Raw264.7 cells were exposed to vehicle (0.1% v/v DMSO) (Veh) or different concentrations (0.5, 2.0 and 10 µmol/l) of BI-9774 (BI) or 10 µmol/l of PF-06409577 (PF) for 16 hours, then cells were lysed and immunoblots were prepared. **d**) Raw264.7 cells were cultured with normal level of glucose (5.5 mmol/l) or low level of glucose (1.0 mmol/l) in the presence of vehicle (0.1% v/v DMSO) (Veh) or 0.5 µmol/l of BI-9774 (BI) for 16 hours. The MIF level in medium was assessed by ELISA. The amount of released MIF was normalized to the total protein amount in each sample. Data are expressed as mean±SEM (**a**-**b**, WT, n=6; D316A, n=5; **c**, n=1; **d**, n=3). Comparisons between groups were made by two-way ANOVA (**a**, **d**) or three-way ANOVA (**b**) with Bonferroni’s multiple comparisons test. **P<0.01; ***P<0.001; ****P<0.0001; ns, not significantly different.

To determine whether AMPK plays a role in MIF release from macrophages, Raw264.7 and mouse primary BMDMs were exposed to PF-06409577 and low glucose (1.0 mmol/l). MIF release was attenuated in the presence of PF-06409577 in both Raw264.7 cells and BMDMs (Fig. 2c, d), and this was replicated by chemically distinct pan-AMPK activator BI-9774 [31] (Fig. 3d). This change in MIF release was not accompanied by a change in cellular MIF expression (ESM Fig. 2).

To examine whether genetic activation of AMPK alters MIF release, BMDMs were obtained from wild type (WT) or AMPKγ1 transgenic mice (hereafter, referred to as AMPK D316A) mice, which display elevated basal AMPK activity as previously described [25]. Phosphorylation of ACC was modestly increased in BMDMs isolated from AMPK D316A mice, although not significantly (Fig. 3a). Consistently, MIF release modestly but not significantly decreased from BMDMs isolated from AMPK D316A (Fig. 3b). PF-06409577 further decreased the MIF release, signficantly, at both normal glucose (5.5 mmol/l) and low glucose (1.0 mmol/l) levels (Fig. 3b).

### 3.3 AMPK activation prevented low glucose-induced changes in cellular metabolism

To examine whether macrophage cellular metabolism was altered following low glucose exposure and the effect of pharmacological activation of AMPK, extracellular flux analyses was performed in Raw264.7 cells and BMDMs at one hour (Fig. 4a, b), three hour (Fig. 4c, d) and 16 hour (Fig. 4e, f) following exposure to reduced glucose levels ± PF-06409577. The ratio of extracellular acidification rate (ECAR) over oxygen consumption rate (OCR) was quantified to measure the contribution of aerobic glycolysis and oxidative phosphorylation to metabolism. Raw264.7 cells and BMDMs exposed to low glucose had a significantly decreased ECAR/OCR ratio at one hour and three hour, which was not apparent at 16 hours. The effect was largely driven by a reduction in ECAR with a more modest increase in OCR at one and three hours. Interestingly pharmacological activation of AMPK by PF-06409577 prevented the low glucose-induced reduction ECAR/OCR ratio to levels comparable to control. Despite the significant reduction in glucose availability, we found that intracellular levels of ATP were well defended over the time course, with no significant changes observed during low glucose exposure or with PF-06409577 treatment (ESM Fig. 4).

**Figure 4.**
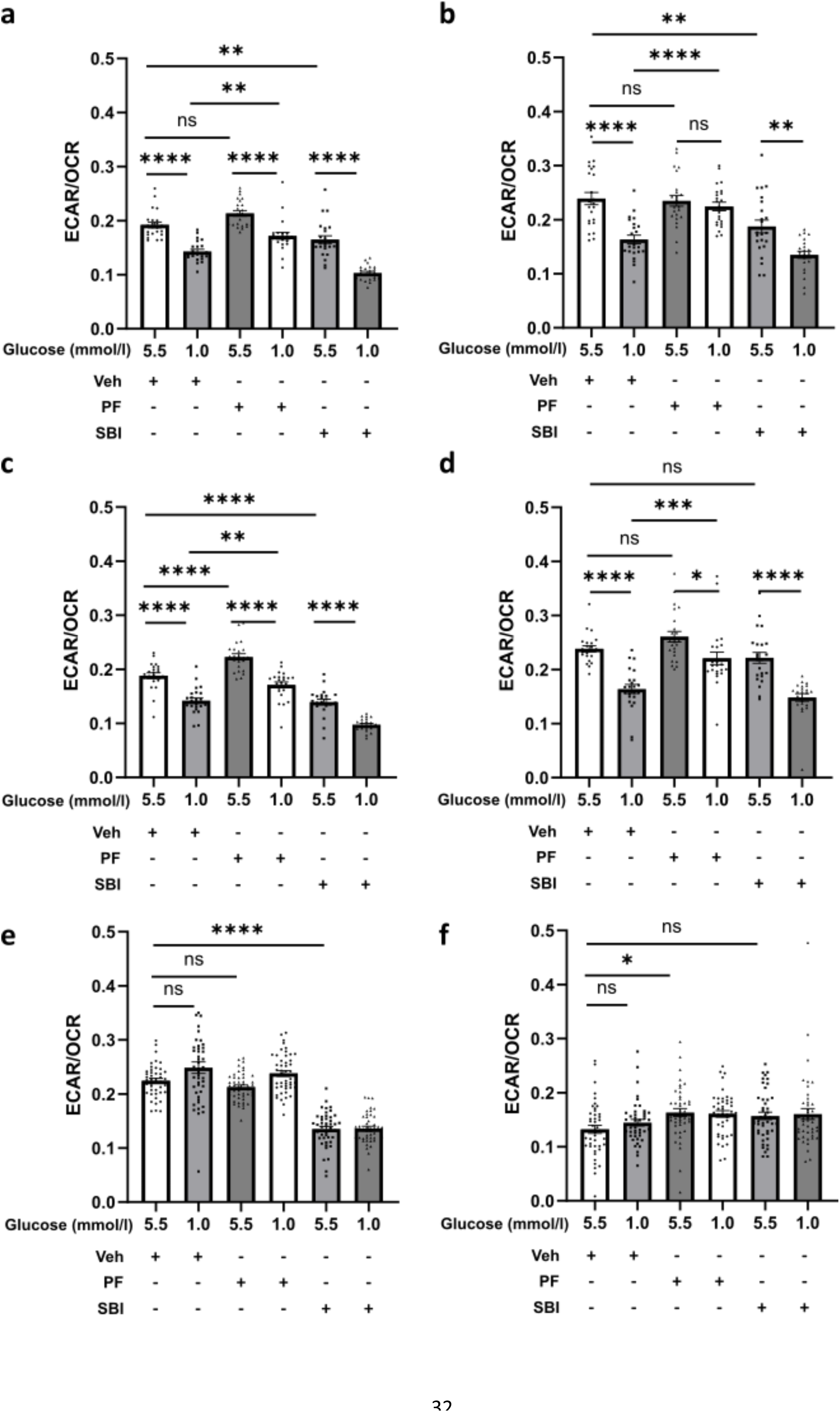
Low glucose reduced the ratio of glycolysis to oxidative phosphorylation, and PF-06409577 restored the ratio in Raw264.7 cells and BMDM at 1 hour and 3 hours but not at 16 hours. **a**) Raw264.7 cells and **b**) BMDM were cultured in the presence of different concentrations of glucose (5.5 mmol/l and 1.0 mol/l) with vehicle (0.1% v/v DMSO) (Veh) or 10 µmol/l of PF-06409577 (PF) or 30 µmol/l of SBI-0206965 (SBI) for 1 hour (**a**, **b**), 3 hours (**c**, **d**), or 16 hours (**e**, **f**) in Seahorse XF96 cell culture microplates. Then the contribution of glycolysis over oxidative phosphorylation was expressed as the ratio of extracellular acidification rate (ECAR) to oxygen consumption rate (OCR) using the Seahorse BioAnalyzer XFe96. Data was expressed as mean± SEM (**a**-**d**, n=23; **e**-**f**, n=45 or 48). Comparisons between groups were made by two-way ANOVA Bonferroni’s multiple comparisons test. *P<0.05; ***P<0.001; ****P<0.0001; ns, not significantly different.

### 3.4 Low glucose induced oxidative stress

We next probed for changes in autophagy given that impairment of autophagy leads to increased oxidative stress [32], and loss of autophagy enhances MIF release from macrophages [33], yet AMPK activation promotes autophagy by phosphorylation ULK1 and mTOR [34]. The ratio of LC3I/II was significantly decreased, indicating autophagy activation in Raw264.7 cells exposed to low glucose (Fig. 5a). However, in contrast to expectations, neither PF-06409577 nor SBI-0206965 altered the conversion of LC3 I to II in both Raw264.7 cells and BMDMs (Fig. 5b, c).

**Figure 5.**
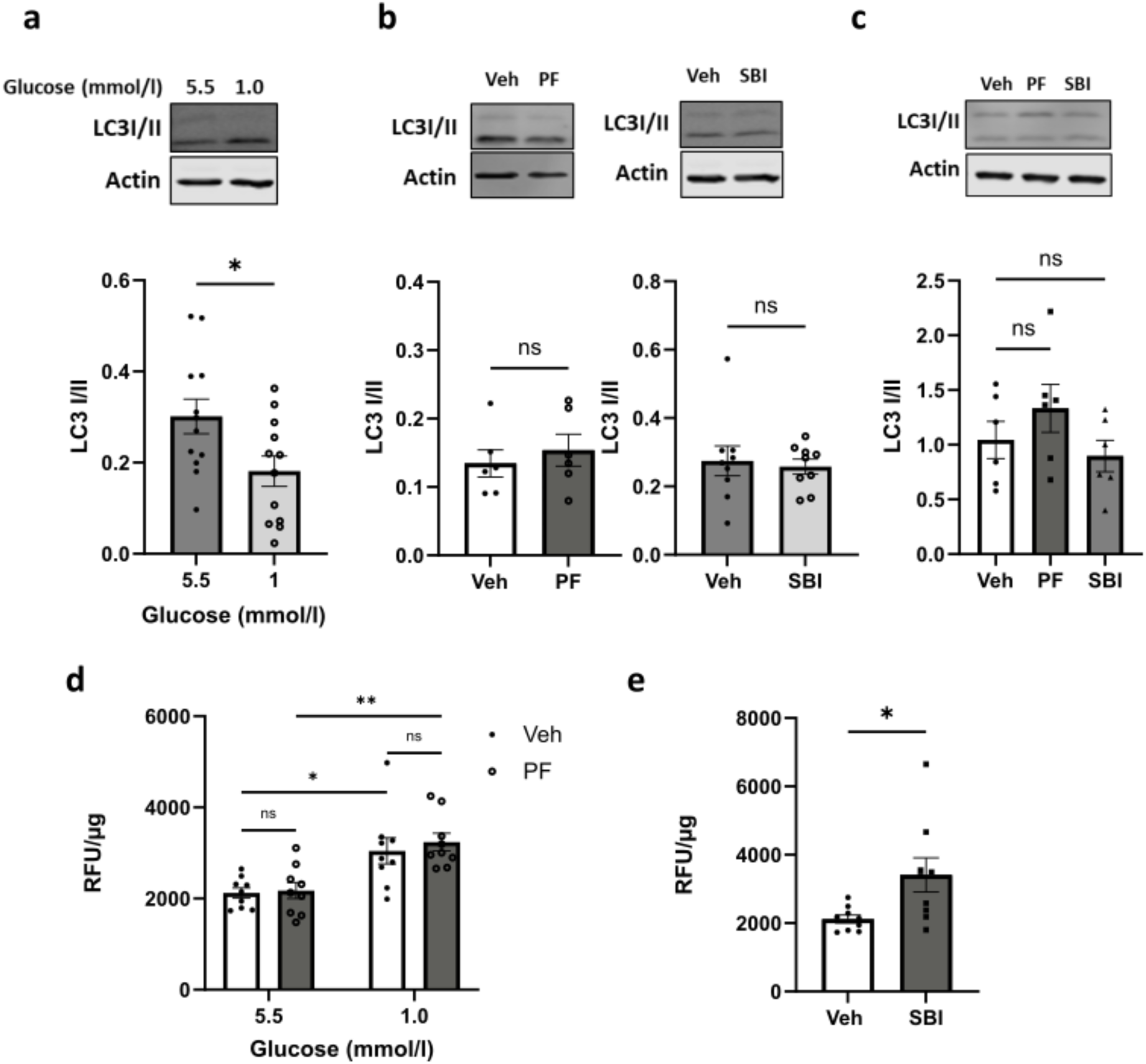
Low glucose activated autophagy and elevated ROS level, but PF-06409577 did not altered autophagy or ROS level in Raw264.7 cells. **a**) Representative immunoblots of Raw264.7 cultured with either 5.5 mmol/l or 1.0 mmol/l glucose for 16 hours. **b**) Representative immunoblots of Raw264.7 cells exposed to vehicle (0.1% v/v DMSO) (Veh) or PF-06409577 (PF) or SBI-0206965 (SBI) for 16 hours. **c**) Representative immunoblots of BMDMs exposed to vehicle (0.1% v/v DMSO) (Veh) or PF-06409577 (PF) or SBI-0206965 (SBI) for 16 hours. Densitometric analysis of immunostaining for LC 3 I and LC3 II. The ratio of LC3 I/II demonstrates the activity of autophagy. **d**) ROS assay results of Raw264.7 cells cultured in the presence of 5.5 mmol/l or 1.0 mmol/l glucose with vehicle (0.1% v/v DMSO) (Veh) or PF-06409577 (PF) for 16 hours. **e**) ROS assay results of Raw264.7 cells exposed to vehicle (0.1% v/v DMSO) (Veh) or SBI-0206965 (SBI) for 16 hours. Following by staining with DCFDA and assessed in Pherastar plate reader and normalized to total protein content in the same well. Data are expressed as mean±SEM (**a**, n=12; **b**, n=6 and 9; **c**, n=6; **d**, n=9; **e**, n=9). Comparisons between groups were made by t test (**a**, **b**, **e**) or by one-way ANOVA (**c**), or by two-way ANOVA with Bonferroni’s multiple comparisons test (**d**). *P<0.05; **P<0.01; ns, not significantly different.

To explore the underlying mechanism, oxidative stress was examined given that reactive oxygen species (ROS) generation has been reported to increase during hypoglycemia [35]. We observed that low glucose significantly elevated ROS levels in Raw264.7 cells (Fig. 5d). Previous studies correlated increased ROS generation with MIF release in cardiomyocytes [36]. Despite the sizable reduction in MIF release following PF-06409577 treatment, this compound did not alter the low glucose-induced ROS production (Fig. 5d). Interestingly SBI-0206965 significantly increased the ROS level (Fig. 5e). However, further examination revealed this to be AMPK-independent, given that the compound increased MIF release from AMPKα1/α2 KO MEF cells (ESM Fig. 5) in cell apoptosis-dependent manner (ESM Fig.7).

### 3.5 Low glucose-induced MIF release was NFĸB dependent

AMPK can inhibit and/or reduce translocation of NFĸB to the nucleus [37], to reduce pro-inflammatory cytokine production. Here TPCA-1 was used to inhibit NFĸB [38], with lipopolysaccharide (LPS) used as a positive control [39]. LPS-induced phosphorylation of NFĸB was blocked by TPCA-1 in Raw264.7 cells, as expected (Fig. 6a) and importantly, TPCA-1 also attenuated low glucose induced MIF release (Fig. 6b), suggesting that NFĸB plays a role in low glucose-induced MIF secretion. There was a modest, non-significant trend towards increased phosphorylated-NFĸB-p65 after low glucose exposure in Raw264.7 cells that was not observed in BMDMs (ESM Fig. 6). Moreover PF-06409577 did not significantly alter NFĸB phosphorylation, however, it did significantly reduce, albeit modestly, the nuclear/cytoplasmic ratio of NFĸB (Fig. 6d)

**Figure 6.**
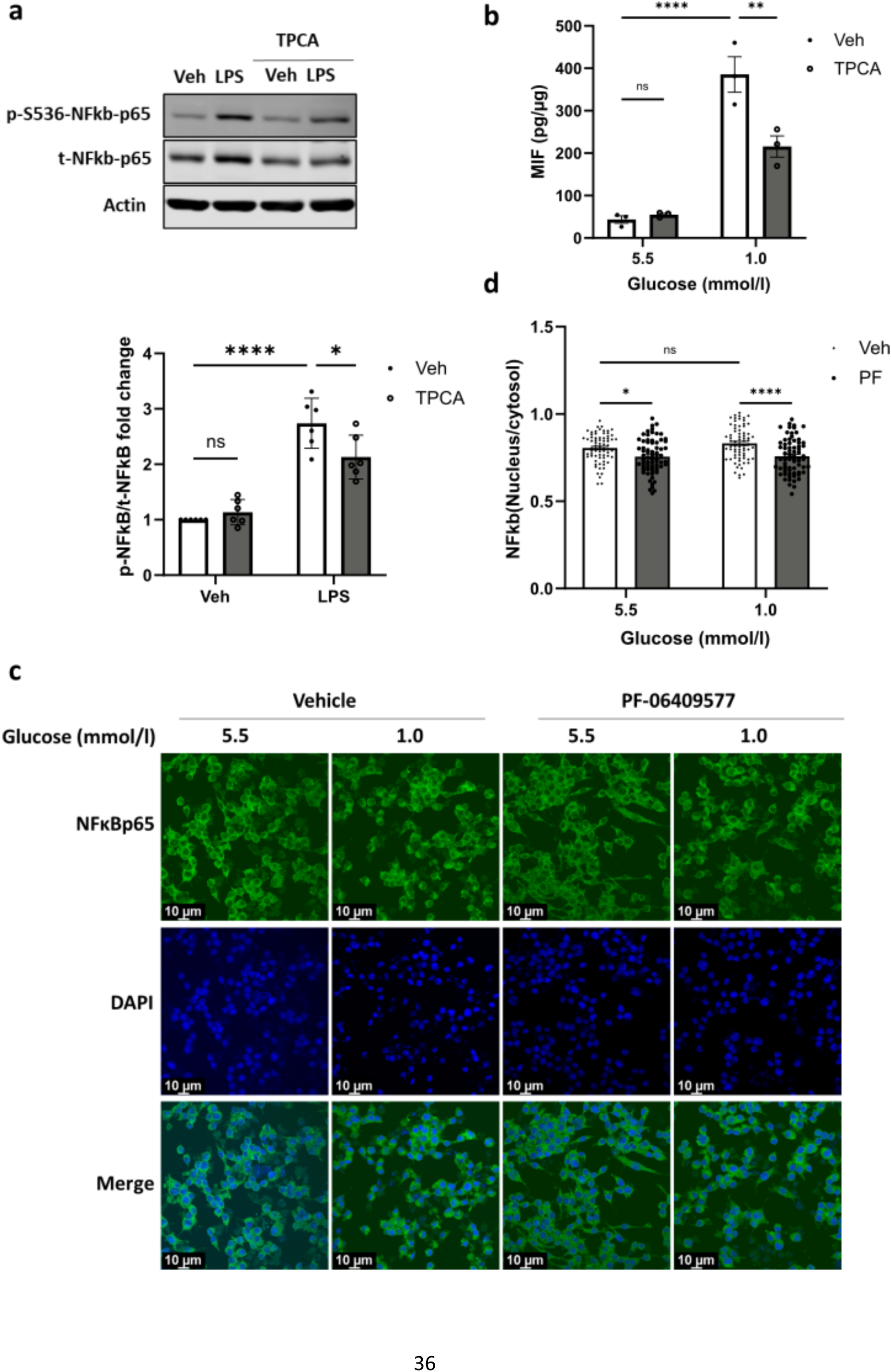
PF-06409577 reduced the translocation of NFĸBp65 and TPCA inhibited LPS induced NFĸB activation and low glucose induced MIF release in Raw264.7 cells. **a)** Representative immunoblots of Raw264.7 cells exposed to vehicle (0.1% v/v DMSO) (Veh) or 100 ng/ml of LPS in the presence of 1 µmol/l of TPCA for 6 hours. Densitometric analysis of immunostaining for phosphorylated protein was normalized to total protein level. p-NFĸB, phospho-S536-NFĸB-p65 (p-S536-NFĸB-p65); t-NFĸB indicates total NFĸB-p65 (t-NFĸB). b) Raw264.7 cells were cultured with either 5.5 mmol/l or 1.0 mmol/l glucose for 16 hours and in the presence of 1 µmol/l of TPCA for the last 6 hours, MIF released in media was assessed in ELISA. c) Raw264.7 cells seeded on coverslips were cultured with either 5.5 mmol/l or 1.0 mmol/l glucose for 16 hours and in the presence of 10 µmol/l of PF-06409577 for 16 hours following by immunocytochemistry staining with NFĸBp65 as Alexa fluor488 and nucleus with DAPI. d) The green fluorescence intensity in the nucleus and cytosol was measured in FIJI and data were collected from three independent experiments, 24 cells from four separate images in each experiment. Data are expressed as mean±SEM (a, n=6; b, n=3; d, n=72). Comparisons between groups were made by two-way ANOVA with Bonferroni’s multiple comparisons test. *P<0.05; **P<0.01; ****P<0.0001; ns, not significantly different.

## 4. Discussion

Our studies demonstrate for the first time low glucose-induced MIF release from primary BMDMs, Raw264.7 cells and BMDM cells isolated from AMPK gain-of-function mice. Previous studies have demonstrated an association between circulating MIF and adipose tissue inflammation, development of insulin resistance and atherosclerosis and β-cell dysfunction [40]. In the present study, we tested the hypothesis that MIF release may be increased by low glucose and attenuated by pharmacological AMPK activation. Other studies have shown that MIF expression was increased in a glucose concentration-dependent manner in cardiomyoblasts and adipose cells [9, 41]. In our study, we observed no change in the cellular MIF expression in macrophages. However consistent with the finding in adipocytes [41] we found that low glucose elevated MIF secretion from macrophages. Currently, the mechanisms of MIF release are poorly defined though cell death and loss of autophagy likely contribute to MIF release [33]. We ruled out cell death in our studies, at least over the time course studied. Moreover, we observed activation of autophagy following low glucose suggesting loss of autophagy is unlikely to mediate increased MIF release shown here. MIF is constitutively expressed and contained within readily-releasable pools in contrast to other pro-inflammatory cytokines and suggests that MIF may act as first responder during hypoglycemia [42]. In contrast to expectations, we did not observe changes in other pro-inflammatory cytokines such as TNFα and IL-1 β release from macrophages (data not shown). It will be interesting to determine in future studies whether low glucose generates a greater and broader increase in pro-inflammatory cytokine production from polarized macrophages, from macrophages isolated from models of diabetes/people with diabetes or during glucose recovery following a hypoglycemia-like challenge. Polarized pro-inflammatory M1 macrophages switch to aerobic glycolysis, and this metabolic switch is associated with reduced AMPK activation [19], whereas anti-inflammatory M2 macrophages utilize mainly oxidative metabolism to generate ATP. In our naïve macrophages, we did not see a metabolic shift towards glycolysis and in contrast, low glucose activated AMPK transiently and decreased the ratio of glycolysis to oxidative phosphorylation. This effect persisted for up to 3 hours, suggesting a successful metabolic adaptation of preserve intracellular ATP levels. Interestingly, pharmacological AMPK activation inhibited or reduced the necessity for this metabolic adaptation to low glucose. This begs the question as to what, if any, is the mechanistic difference between physiological (low glucose) and pharmacological AMPK activation? We noted modest increases in AMPK pathway activation in response to low glucose that coincided with increased MIF release. We tried to examine whether AMPK inhibition altered MIF release, however, SBI-0206595 increased MIF release in an AMPK-independent manner, mostly likely by increasing ROS production. BMDMs isolated from AMPK D316A allowed us to examine non-pharmacological effects. In line with the pharmacology studies, modest genetic AMPK activation attenuated low glucose-induced MIF release. These data lend support to the idea increasing levels of AMPK activity produce greater suppression of MIF release and that the regulation of MIF by AMPK is most apparent when there is a stress to work against.

AMPK is generally regarded as a negative regulator of pro-inflammatory cascades, although the underlying mechanism is still unclear [43]. Hematopoietic AMPK β1-containing AMPK complexes reduce mouse adipose tissue macrophage inflammation and insulin resistance in obesity [44]. Our data demonstrate that pharmacological AMPK activation, using two different AMPK activators, attenuates MIF release from macrophages. Previous studies have shown that hypoglycemia/low glucose induces oxidative stress [14], and separate studies have shown that AMPK can inhibit oxidative stress [45] by activation of autophagy to scavenge ROS [46]. There is also evidence that oxidative stress induces MIF release [47]. Accordingly we demonstrate that low glucose mediated oxidative stress may drive MIF release. However, PF-06409577 did not alter low glucose-induced ROS production, suggesting that AMPK-mediated suppression of MIF occurs further downstream or that increased ROS is a bystander event.

Surprisingly, we found no evidence that PF-06409577 altered autophagy. It has been suggested that AMPK regulation of autophagy is a dual ‘fail-safe’ mechanism mediated by inactivating of TORC1, the master regulator of autophagy, and direct phosphorylation of ULK1, an initiator of autophagy [48]. Despite a recent report demonstrating PF-06409577 mediated reductions in the ratio of LC3 I/II in osteosarcoma cells [49], this was not observed in macrophages even at higher concentrations (10 µmol/l vs. 1 µmol/l). PF-06409577 did not regulate autophagy in our study, at least under the conditions tested. Collectively, our findings suggest that low glucose induced MIF release is likely mediated, at least in part by increasing oxidative stress and that AMPK acts downstream of oxidative stress to suppress MIF release. The regulation of MIF secretion is poorly understood. Unlike most other cytokines, MIF lacks the N-terminal signal peptide that guides secretion through the conventional endoplasmic reticulum (ER)-Golgi route, therefore is secreted via an unconventional pathway [50]. Previous studies demonstrated that inhibition of autophagy using RNAi or autophagy inhibitor, 3-methyladenine (3-MA) enhanced MIF release from macrophages [33]. Consistently, we observed that AMPK/ULK/autophagy inhibitor SBI-0206965 substantially enhanced MIF release from macrophages, despite no effect on LC3 conversion. In the present study, low glucose-induced autophagy coincided with increased MIF release, indicating that autophagy is not directly involved in the regulation of MIF secretion from macrophages. Here, it is likely that suppression of MIF occurs through an AMPK-NFĸB signalling pathway given the role of NFĸB in the anti-inflammatory actions of AMPK in other pathological states [51]. AMPK inhibits the translocation of NFĸB where it can otherwise activate transcription of genes involved in pro-inflammatory signalling [39] and we confirmed this in the present study, although this effect was modest. Similarly, NFĸB inhibition attenuated low glucose-induced MIF secretion, further demonstrating that NFĸB is involved in MIF release. Interestingly, MIF stimulates AMPK activation through CD74, promotes glucose uptake and protects the heart during ischaemia-reperfusion injury in cardiomyocytes [52] and increases glucose utilisation in skeletal muscle [53]. In our study, we did not observe significant AMPK activation when MIF release was enhanced after low glucose exposure (i.e. at 16 hrs). It is possible that the autocrine-paracrine effect of MIF on the AMPK activation is too modest or transient to be detected in our conditions. According to studies in other cells, induction of significant AMPK activation by MIF requires concentrations of greater than 100 ng/ml, which is greater than the highest MIF concentrations detected here [54, 55]. However, given that we did not find production of other pro-inflammatory cytokines in our studies, it is plausible that MIF is acting in a protective and or compensatory manner to help cells defend against low energy availability.

## 5. Conclusion

In summary, these data suggest that macrophage AMPK acts to limit MIF release under pathological states where increased ROS-mediated activation of NFĸB occurs. Moreover, pharmacological AMPK activation suppresses MIF release without altering the generation of ROS, at least in response to low glucose in the current study. Taken together, this suggests that pharmacological AMPK activation could be a strategy for reducing macrophage MIF release in diseases such as diabetes, where glucose variation frequently occur although further study is required to determine the nature of MIF signalling in the context of low glucose reported here.

## Acknowledgements

The authors wish to thank Dr Ana Cruz, Dr Paul Weightman Potter, Dr Katie Partridge, Katherine Pye, Wyn Firth and Asmaa Alkhalidi for technical assistance and insightful comments during the project. The graphical abstract was created with BioRender.com.

## Data Availability

All data generated or analysed during this study are included in the published article (and its ESM). The file is available from the corresponding author upon reasonable request.

---

**Supplementary Figure 1.**
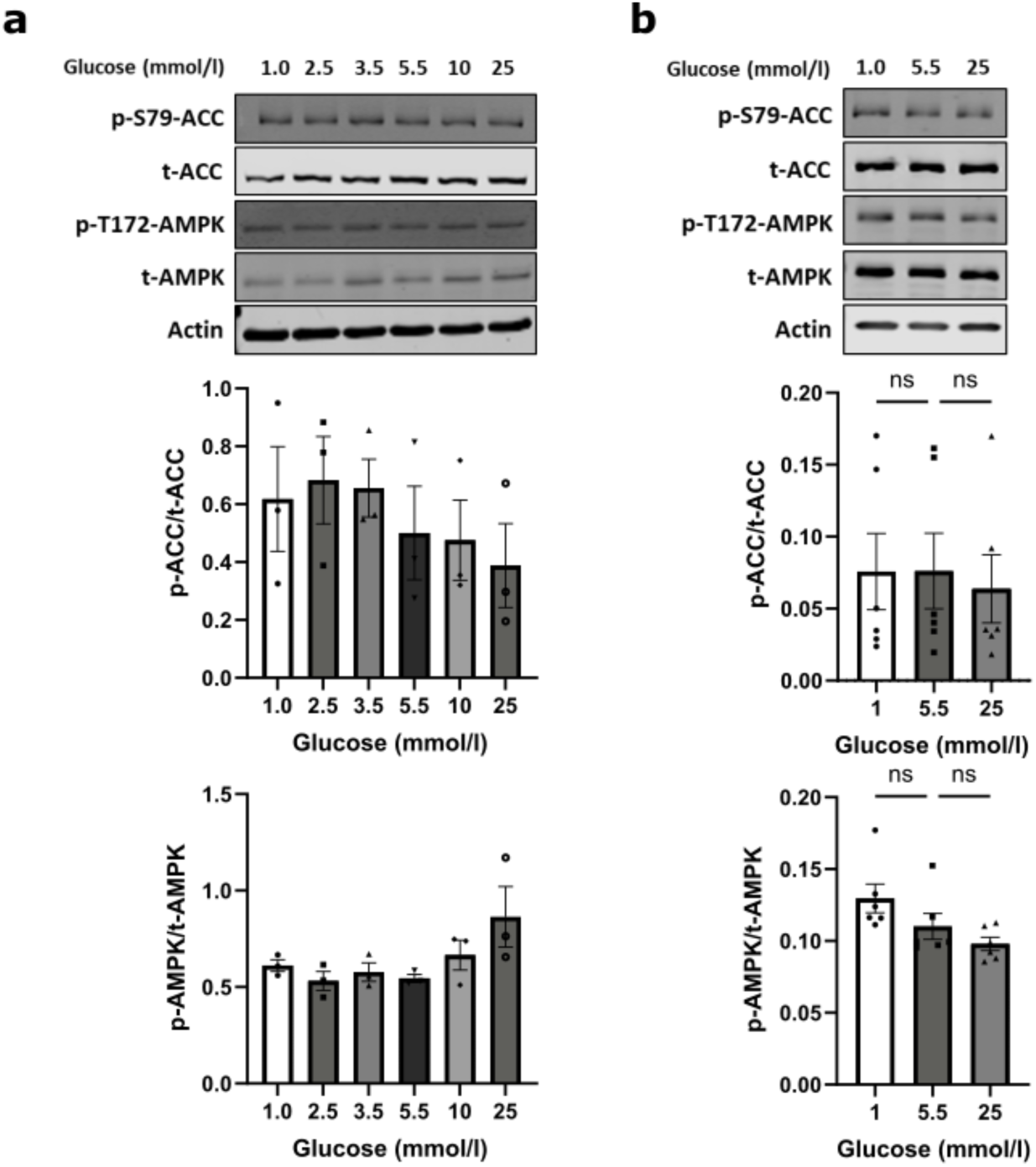
The effect of different concentrations of glucose on AMPK activation in Raw264.7 cells and BMDMs. **a)** Raw254.7 cells and b) BMDMs were cultured with different concentrations of glucose for 16 hours. Cells were then lysed, and immunoblots were prepared. Densitometric analysis of immunostaining for phosphorylated protein was normalised to total protein level. p-ACC, phospho-acetyl-CoA caboxylase (S79) (p-S79-ACC); p-AMPK, phospho-AMP-activated protein kinase (T172)(p-T172-AMPK); t-ACC indicates total acetyl-CoA carboxylase; t-AMPK, total AMP-activated protein kinase. Data are expressed as mean±SEM (a, n=3; b, n=6). Comparisons between groups were made by one-way ANOVA with Bonferroni’s multiple comparisons test, ns, not significantly differnt.

**Supplementary Figure 2.**
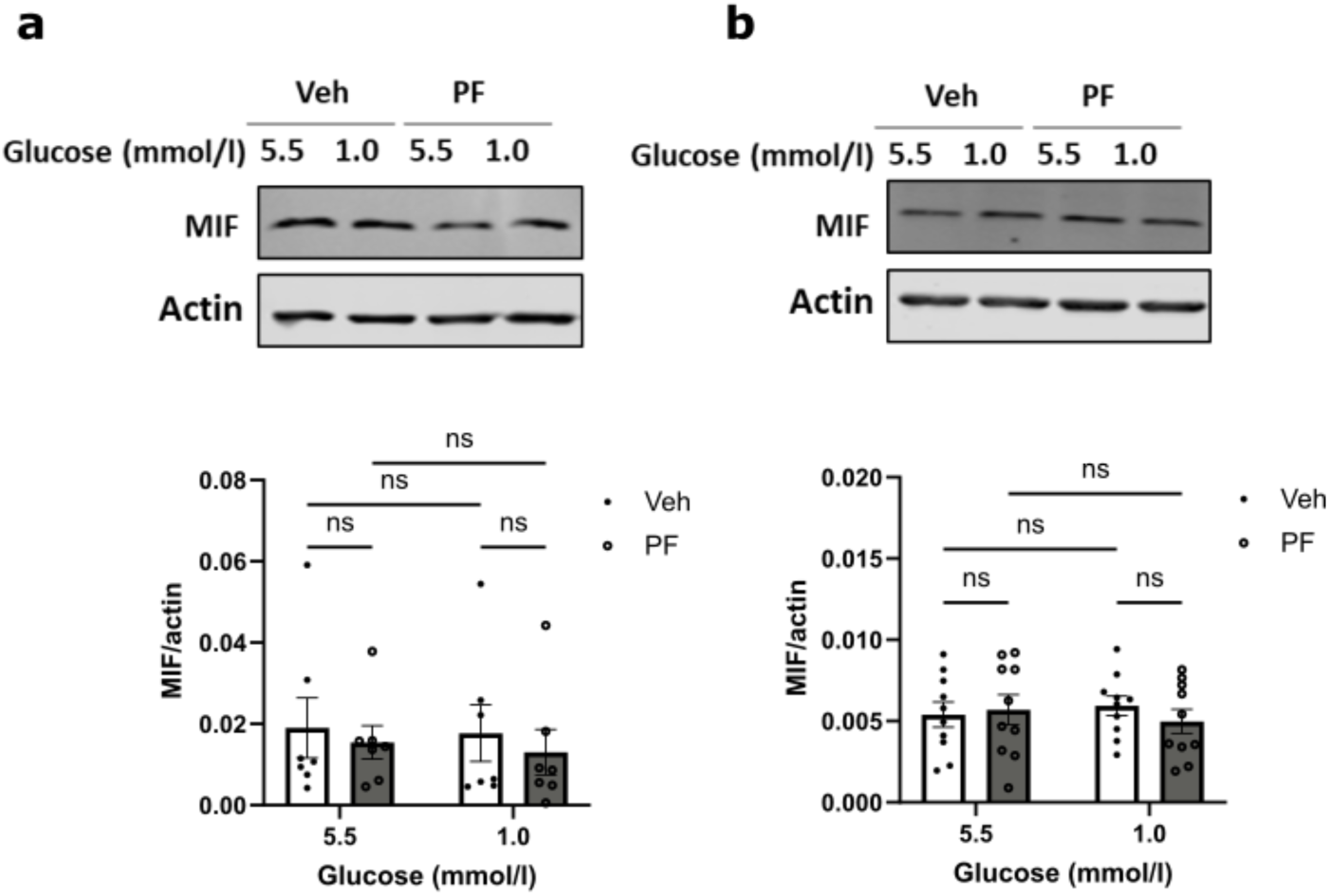
low glucose and/or PF-06409577 did not alter the intracellular expression of MIF in macrophages. **a)** Raw264.7 cells and **b)** BMDMs were cultured with either 5.5 mmol/l or 1.0 mmol/l of glucose in the presence of vehicle (0.1% v/v DMSO) (Veh) or 10 pmol/l of PF-06409577 (PF) for 16 hours. Cells were then lysed, and immunoblots were prepared. Densitometric analysis of immunostaining for MIF was normalised to actin, Data are expressed as mean±SEM **(a,** n=7; **b,** n = 10). Comparisons between groups were made by two-way ANOVA test, ns, not significantly different.

**Supplementary Figure 3.**
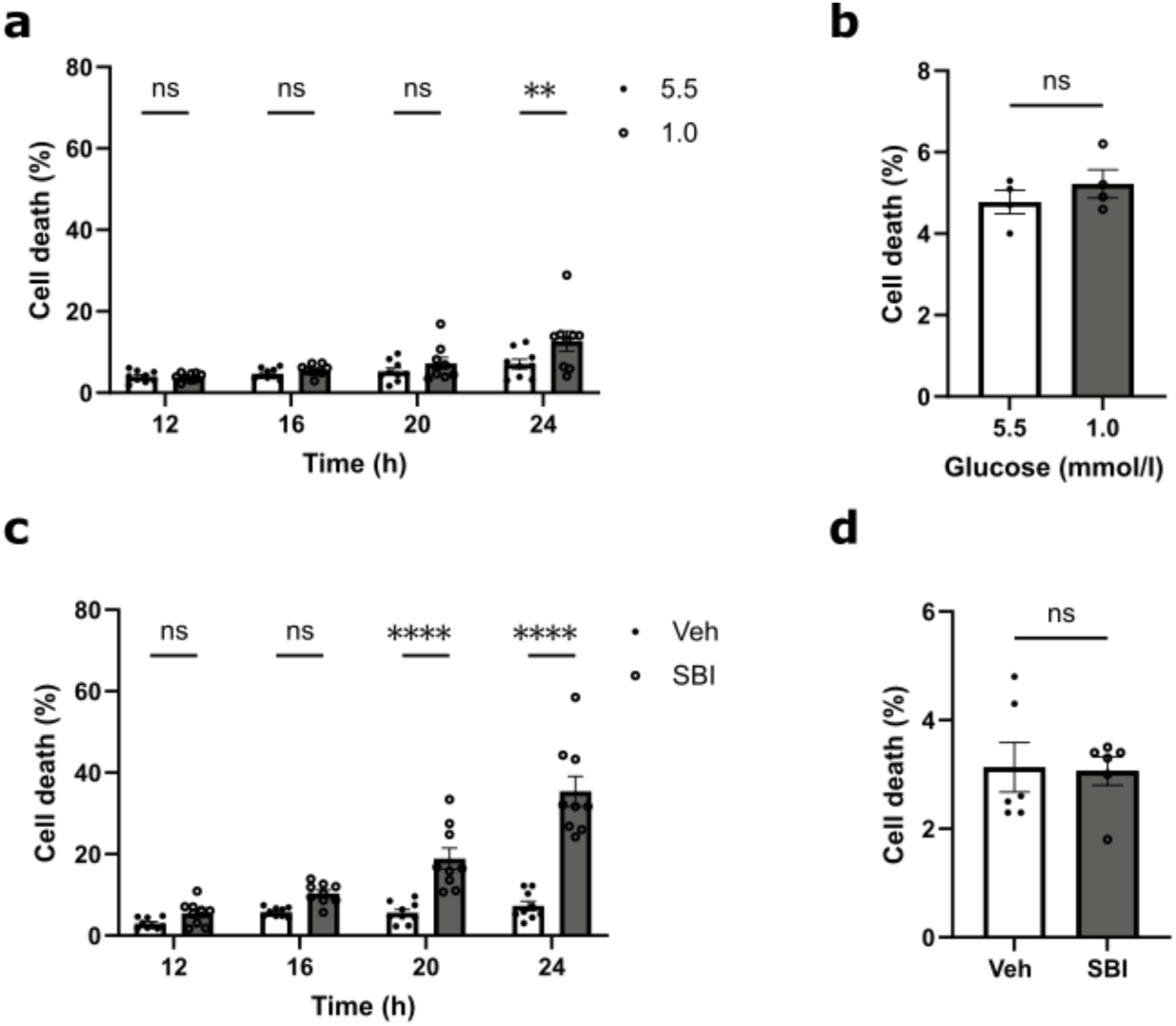
Low level of glucose (1.0 mmol/l) and SBI-0206965 increased cell death over time in Raw264.7 cells, but no significant cell death were detected at 16 hours for either Raw264.7 cells or BMDMs. **a)** The time course of cell death of Raw264.7 cells cultured with 5.5 mmol/l or 1.0 mmol/l glucose for up to 24 hours, b) The cell death of BMDMs exposed to 5.5 mmol/l or 1.0 mmol/l glucose for 16 hours, c) The time course of cell death of Raw264.7 cells exposed to vehicle (0.1% v/v DMSO) (Veh) or 30 pmol/l of SBI-0206965 (SBI) for up to 24 hours, d) The cell death of BMDMs exposed vehicle (0.1% v/v DMSO) (Veh) or 30 pmol/l of SBI-0206965 (SBI) for 16 hours. Cells viability was assessed at indicated time points by staining with propidium iodide followed by flowcytometry. The numbers of postive staining cells were expressed as percentage of total cell numbers. Data are expressed as mean±SEM (a, n=9; b, n=3; c, n=9; d, n=6), Comparisons between groups were made by two-way ANOVA with Bonferroni’s multiple comparisons test (a, c) or unpaired t test (b, d). **P<0.01; ns, not significantly different.

**Supplementary Figure 4.**
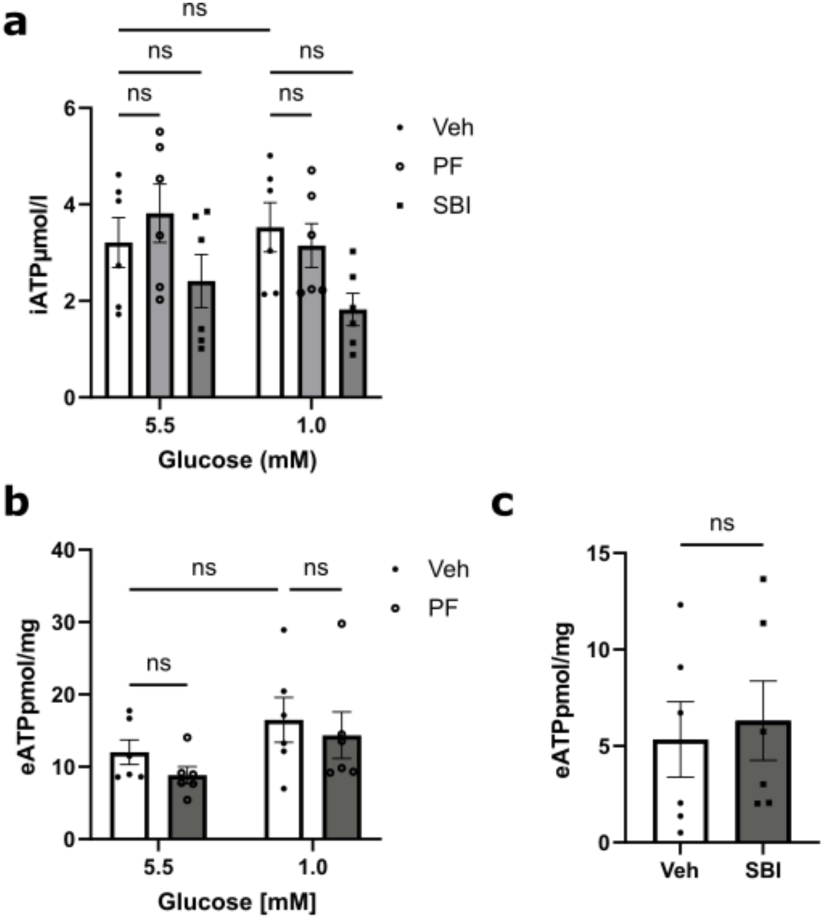
PF-06409577 or SBI-0206965 eATP in Raw264.7 cells. **a)** Raw264.7 cells were cultured did not alter the (1.0 mmol/l), levels of iATP or in the presence of 5.5 mmol/l or 1.0 mmol/l of glucose with vehicle (0.1% v/v DMSO) (Veh) or 10 μmol/l of PF-06409577 (PF) or 30 pmol/l of SBI-0206965 (SBI) for 16 hours. The cells in the well were then lysed and intracellular ATP (iATP) were measured using luminescence based assay ATPIite. **b)** The extracellular ATP (eATP) released in the media was measured using luminescence based assay ATPIite and normalised to total protein content in the same dishes **(b, c).** Data are expressed as mean± SEM (n=6), Comparisons between groups were made by two-way ANOVA with Bonferroni’s comparison test **(a, b),** or unpaired t test **(c).** ns, not significantly different.

**Supplementary Figure 5.**
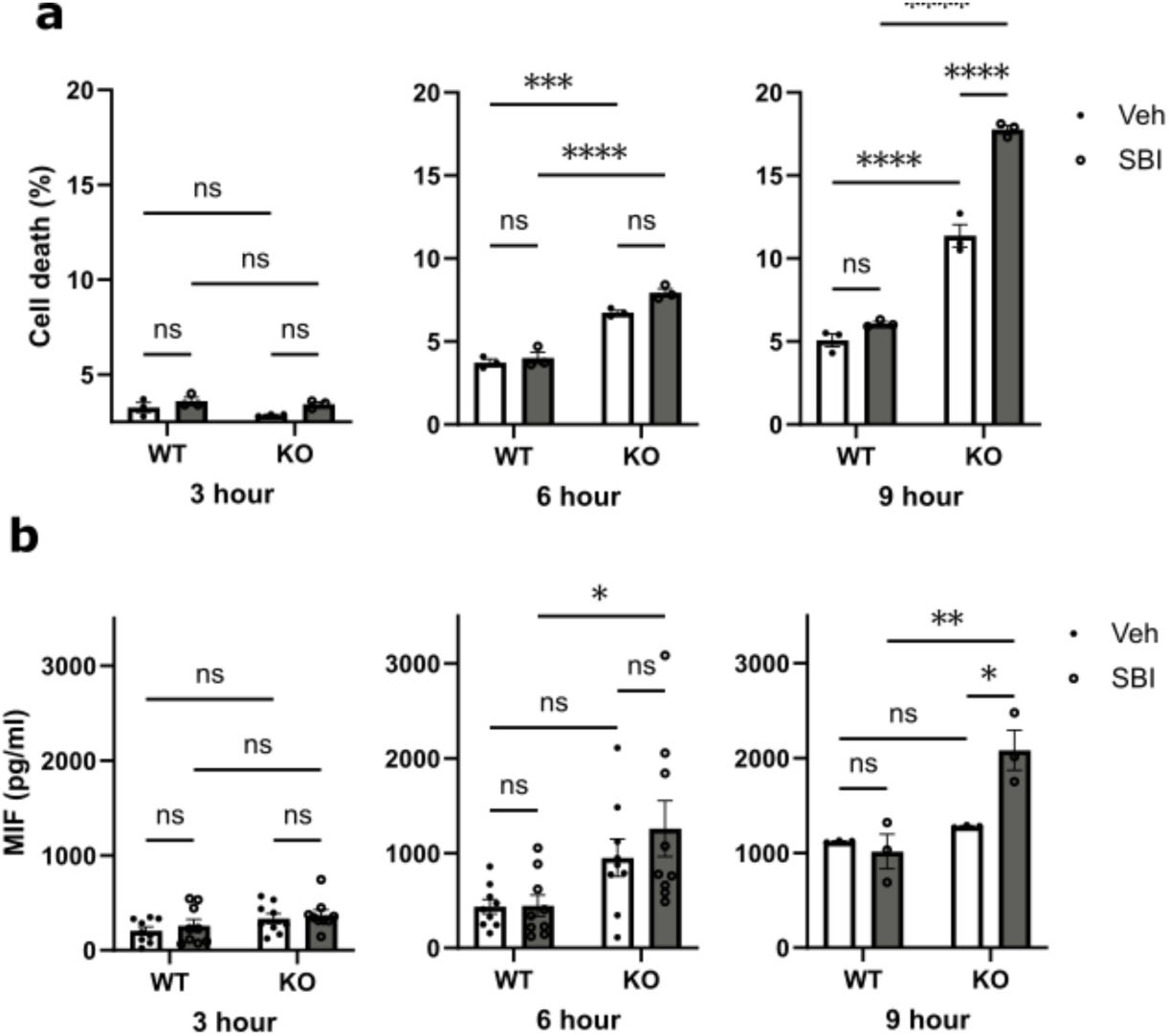
The cell death of AMPK a1/2 (-/-) MEFs was elevated after 6 hours and SBI-0206965 induced MIF release and cell death from AMPK a1/2 (-A) MEFs at 9 hours. **a)** Wild-type (WT) and AMPK a1/2 (-/-) knock out (KO) MEF cells wer were cultured with vehicle (0.02% v/v DMSO) (Veh), or 10 pmol/l of SBI-0206965 (SBI) for 3 hours, 6 hours, and 9 hours. Cells viability was assessed by staining with propidium iodide followed by flowcytometry. The numbers of postive staining cells were expressed as percentage of total cell numbers, b) The MIF levels in medium was assessed by ELISA. Data are expressed as mean±SEM (a, n=3; b, n=9 for 3 and 6 h, n=3 for 9 h). Comparisons between groups were made by two-way ANOVA with Bonferroni’s multiple comparisons test. *P<0.05: **P<0.01; ***P<0.001: ****P<0.0001; ns, not significantly different.

**Supplementary Figure 6.**
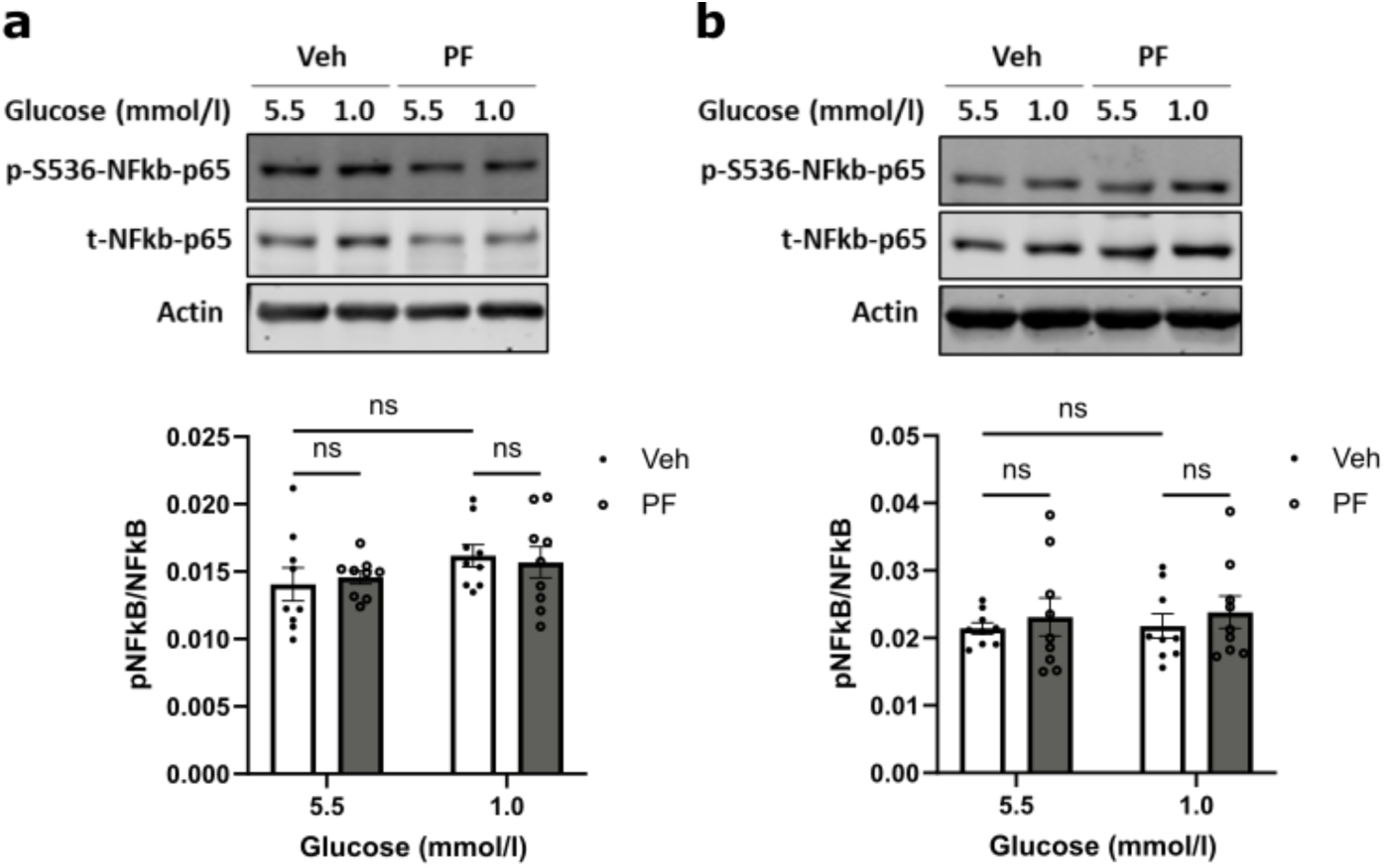
Low glucose or PF-06409577 did not alter the level of phosphorylation of NFKB-p65 in Raw264.7 cells or BMDMs. **a)** Raw 264.7 cells or b) BMDM cells were cultured with either 5.5 mmol/l or 1,0 mmol/l of glucose in the presence of vehicle (0.1% v/v DMSO) (Veh) or 10 μmol/l of PF-06409577 (PF) for 16 hours. Cells were then lysed, and immunoblots were prepared. Densitometric analysis of immunostaining for phosphorylated protein was normalised to total protein level, p-S536-NFKB-p65 indicates phosphoylated NFi<B-p65 at 5536; t-hFkB-p65 Indicates total NFKB-p65. Data are expressed as mean±SEM **(a,** n=9; **b,** n = 9). Comparisons between groups were made by two-way ANOVA with Bonferroni’s multiple comparisons test ns, not significantly different.

**Supplementary Figure 7.**
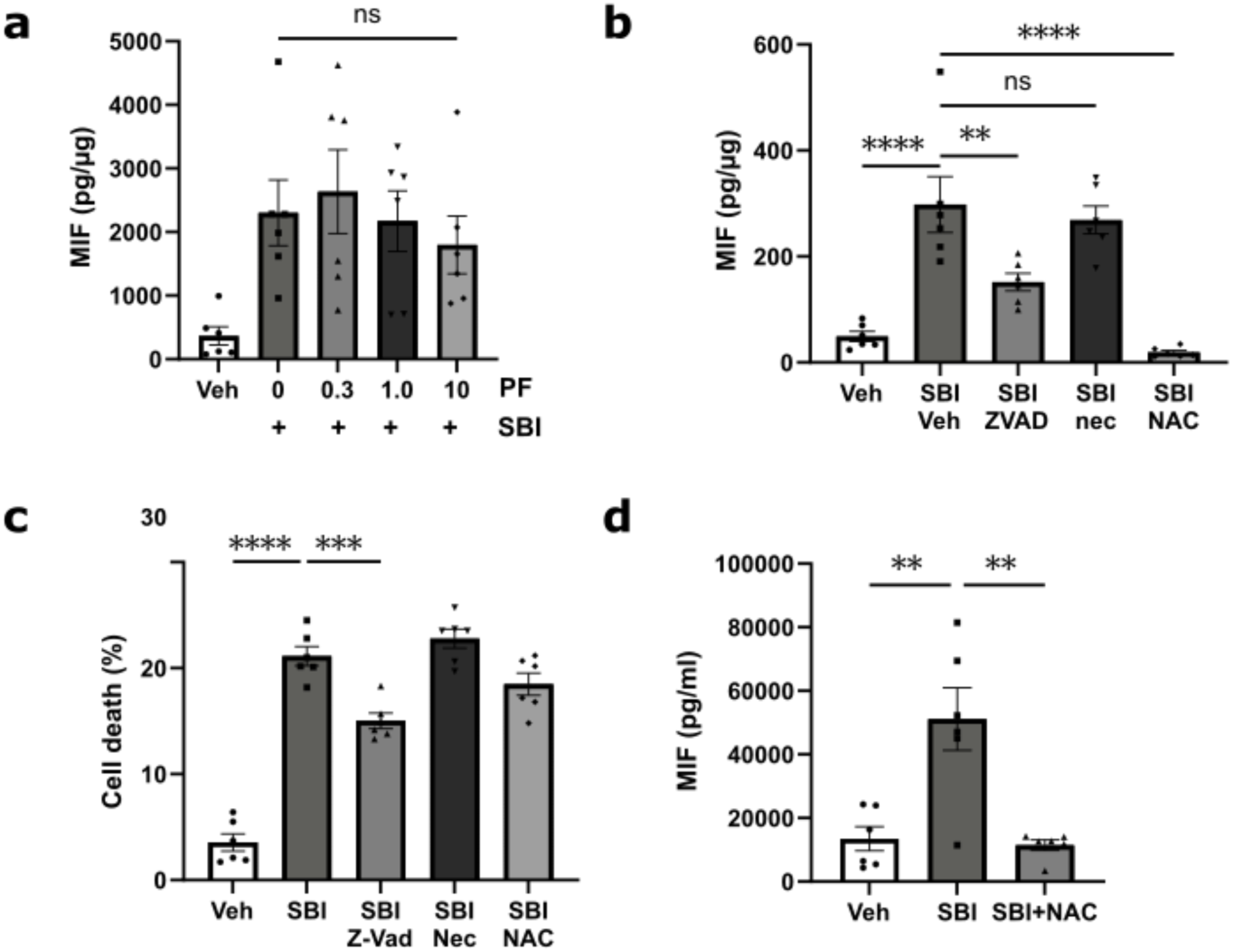
SBI-0206965 increased MIF release and cell death, which was attenuated by Z-VAD but the enhancement of MIF release was not altered by PF-06409577 in Raw264.7 cells. **a)** Raw264.7 cells were cultured with vehicle (0.1% v/v DMSO) (Veh) or 30 pmol/l of SBI-0206965 (SBI) for 15 minutes following by addition of different concentrations (0, 0.3,1.0,10 pmol/l) of PF06409577 (PF) for 16 hours, b) Raw264.7 cells were cultured withh vehicle (0.1% v/v DMSO) (Veh) or 30 pmol/l of SBI-0206965 (SBI) alone, or in the presence of Z-Vad (40 pmol/l), or necrostatin 1(30 pmol/l) (Nec), or NAC (10 mmol/l) for 20 hours. The MIF level in medium was assessed by ELISA and the amount of released MIF was normalised to the total protein amount in each sample c) Cell viability was assessed by staining with propidium iodide followed by flowcytometry. The numbers of positive staining cells were expressed as percentage of total cell numbers, d) NAC (10 mmol/l) was added directly into the SBI-0206965 treated medium before performing ELISA, MIF in the medium was shown in concentration (pg/ml). Data are expressed as mean±SEM (n=6). Comparisons between groups were made by one-way ANOVA with Bonferroni’s multiple comparison test. ***p<0.001; ****P<0.0001; ns, not significantly different.

